# Collagen stiffness modulates MDA-MB231 cell metabolism through adhesion-mediated contractility

**DOI:** 10.1101/272948

**Authors:** Emma J. Mah, Gabrielle E. McGahey, Albert F. Yee, Michelle A. Digman

## Abstract

**Abstract Figure:**
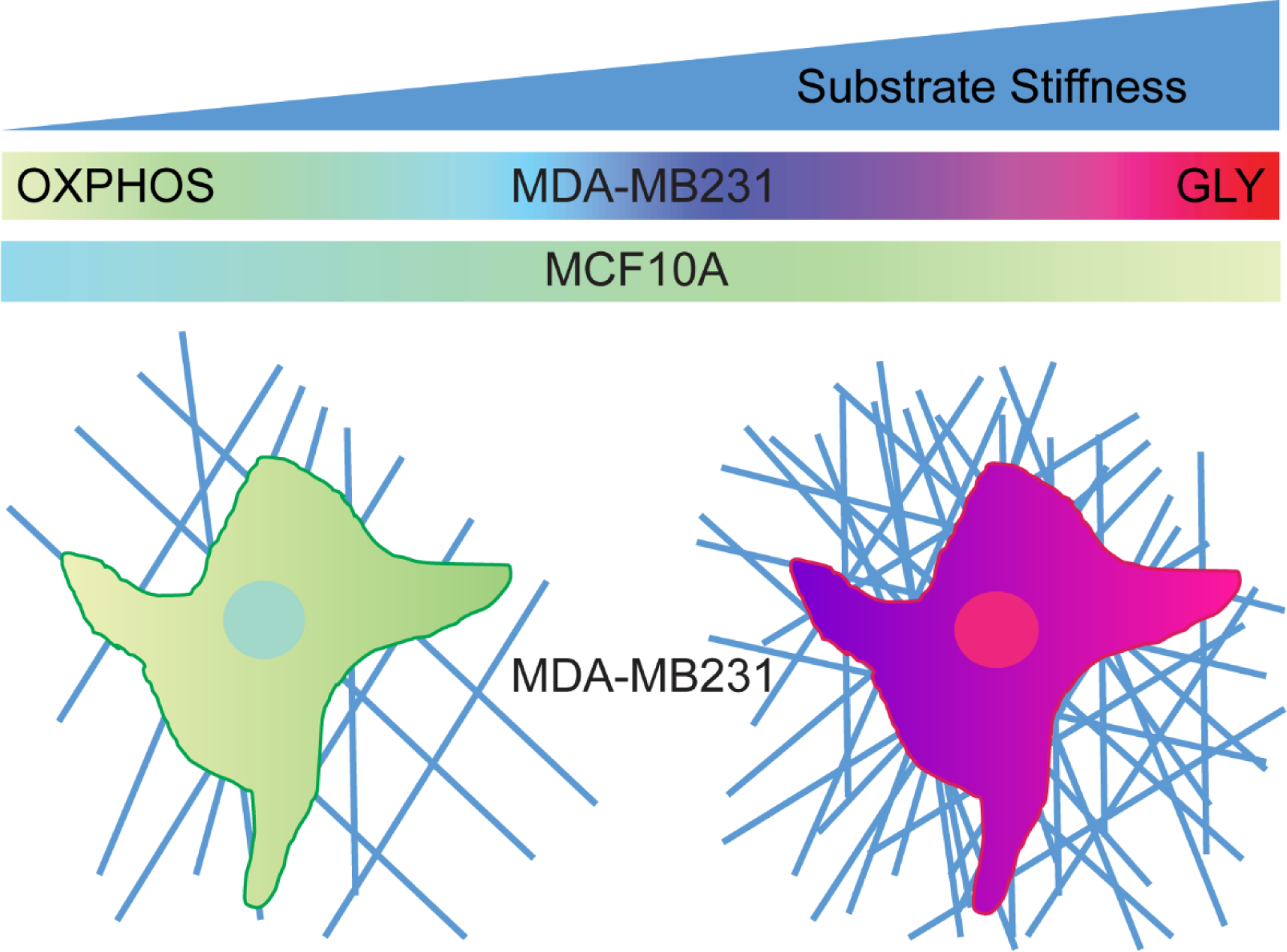
Increasing collagen stiffness causes a shift in highly invasive MDA-MB231 cancer cells from glycolysis to oxidative phosphorylation. Glioma U251MG cells show an opposite trend and MCF10A non-tumorigenic cells have little change in metabolism signatures in response to substrates stiffness.

## Summary

Extracellular matrix (ECM) mechanical properties play a key role in cancer cell aggressiveness. Increasing substrate stiffness upregulates cancer invasion, cell contractility and focal adhesion formation. In addition to matrix properties, alteration in energy metabolism is a known characteristic of cancer cells (i.e., Warburg effect) and modulates cell invasion. However, there has been little evidence to show that substrate stiffness is able to affect cancer cell metabolism. Thus, we investigated changes in energy metabolism in response to varying collagen matrix stiffness in different cancer cells, MDA-MB231, AA375MM and U251MG and non-tumorigenic breast cell line MCF10A. Using the phasor approach to fluorescent lifetime imaging microscopy (FLIM), we measured the lifetime ratio of the free:bound state of NADH and determined if these cells altered their metabolism when plated on varying ECM density. This approach is a powerful tool that allows us to map the metabolic trajectory of each living cell within its cellular compartments. In our studies, we found that MDA-MB231 cells had an increase in bound NADH, indicating oxidative phosphorylation (OXPHOS), as collagen substrate density decreased. When inhibiting myosin-II contractility with Y-27632 or blebbistatin, the MDA-MB231 cells on glass shifted from glycolysis (GLY) to OXPHOS, confirming the intricate relationship between mechanosensing and metabolism in these highly invasive tumor cells. The human glioblastoma cell line, U251MG, showed an opposite trend compared to the invasive MDA-MB231 cells. However, the human melanoma cell line, A375MM did not show any significant changes in metabolic indices when they were grown on surfaces with varying collagen density but changed when grown on glass surfaces. MCF10A cells showed no changes in metabolism across all surfaces. In addition, OXPHOS or GLY inhibitors to MDA-MB231 cells showed dramatic shifts from OXPHOS to GLY or *vice versa*. There were slight changes detected in MCF10A cells. These results provide an important link between cellular metabolism, contractility and ECM stiffness in human breast cancer.

## Introduction

Cancer cells can modulate their energy metabolism to meet nutritional, biosnynthesis and respiration requirements for maintaining malignancy. One of these factors is the metabolic state of the cancer cell due to their tendency to undergo aerobic glycolysis, known as the Warburg Effect (Vander Heiden *et al*., 2009; Liberti and Locasale, 2016; Warburg *et al*., 1927). Although it produces less ATP per molecule of glucose, glycolysis (GLY) is a more rapid way of producing ATP and is able to meet the high demands of energy to fuel processes such as invasion, migration, and matrix degradation (Caino *et al*., 2015; Cunniff *et al*., 2016; Desai *et al*., 2013; Zhao *et al*., 2013). Along with high turnover of ATP production, a byproduct of lactic acid and high acidification also has been shown to benefit cancer cell survival and upregulate invasiveness (Estrella *et al*., 2013).

Mechanical properties of the extracellular matrix (ECM) is also a known factor that regulates cell migration and cancer invasion (Alexander *et al*., 2008; Artym *et al*., 2015; Paszek *et al*., 2005; Seewaldt, 2014; Wells, 2008). Cells interact with the surrounding ECM through integrin-mediated adhesions and focal adhesions (FAs), that are clusters of over 150 proteins (Kanchanawong *et al*., 2010; Liu *et al*., 2015). These complexes tether to the cell’s mechanosensing network through actin filaments and regulate processes such as adhesion, migration, and proliferation (Bugyi and Carlier, 2010; Gardel *et al*., 2010; Hirata *et al*., 2014; Ponti *et al*., 2004). Recent studies have shown that integrin-mediated adhesion interact with the metabolic pathway of the cell through the PI3K/AKT/mTOR pathway and that this could be a potential method of switching the Warburg effect (Ata and Antonescu, 2017; Levental *et al*., 2009; Yang *et al*., 2015). Many of these studies use biochemical assays which are invasive and often lose information which exist in live cell samples. In our approach, we used a non-invasive fluorescent imaging technique to measure ECM density, study live cell behavior and map energy metabolism within each cell.

Fluorescent lifetime imaging microscopy (FLIM) has been shown to be a powerful technique to measure metabolic indices of live-cells (Bird *et al*., 2005; Cinco *et al*., 2016; Datta *et al*., 2015; Ma *et al*., 2016; Provenzano *et al*., 2009; Sameni *et al*., 2016; Stringari *et al*., 2012). By looking at the fluorescent lifetime of nicotinamide adenine dinucleotide (NADH), a metabolite involved in OXPHOS and GLY, we can determine the population of free and bound NADH due to their difference in lifetime decay. This will allow us to quantify the “metabolic trajectory”, known as the “M trajectory”, of the cell at every pixel of our image and determine if the cell is undergoing OXPHOS or GLY (Stringari *et al*., 2012). The advantage of this imaging technique is that it is non-invasive and is able to image real-time changes in metabolism.

For this study, we measured free and bound populations of NADH within different cancer cell lines and a non-tumorigenic cell line when seeded on collagen substrates of different concentrations (1.2 mg/mL and 3.0 mg/mL) and on glass. The microstructural properties of this substrate, including collagen density and fiber diameter, were measured using image correlation spectroscopy (Raub *et al*., 2008). The 3.0 mg/mL and 1.2 mg/mL collagen substrate had collagen fibers of similar size, but the 3.0 mg/mL collagen substrate gave rise to a denser ECM than the 1.0 mg/mL substrate by 3X. The 3.0 mg/mL collagen substrate was also determined by rheology to be one order of magnitude stiffer than the 1.2 mg/mL collagen substrate. The highly metastatic breast cancer cell line MDA-MB21 showed a shift towards a more glycolytic signature as substrate stiffness increased. Inhibition of cell contractility with addition of Y-27632 shifted all the cells on all substrates to a more OXPHOS signature compared to their uninhibited controls. In addition, we observed more stable focal adhesions on our stiffer substrates through raster image correlation analysis by measuring the diffusion of talin-GFP, focal adhesion protein which binds to integrin and the actin cytoskeleton structure. This further shows that integrin mediated adhesions behave as mechanosensors and these adhesions can alter metabolism.

Non-tumorigenic breast cell line MCF10A showed no significant changes in NADH free:bound ratio across all surfaces, indicating that this property only appears in MDA-MB231 cell lines. Other cancer cell types, U251MG glioma and A375MM melanoma cell lines, were evaluated under the same conditions. The U251MG cells had an opposite trend of NADH free:bound ratio signatures across the substrates compared to MDA-MB231 cells. A375MM cells did not adhere well to the collagen substrates which caused them to have no significant difference in their NADH free:bound ratio; which indicates that the mechanosensing network must be established in order to undergo metabolic reprogramming. Inhibition of OXPHOS or GLY in MDA-MB231 cells showed shifts in NADH free:bound ratio with respect to each treatment towards their metabolic counterparts across all surfaces, and further confirmed that it is indeed the metabolism that is being altered by the ECM. MCF10A cells showed a shift when OXPHOS was inhibited only on our denser collagen and on glass substrates when GLY was inhibited. The results found in our work here show that both the mechanosensing and metabolism pathways are interconnected and can be modulated through ECM mechanical properties. This will provide further information to develop cancer therapies which target either or both of these pathways to decrease cancer cell invasion.

## Results

### Collagen characterization measurements

Tilghman *et al.* postulated that cellular metabolism can be altered when MDA-MB231 cells are cultured on soft (300 Pa) versus stiff (19200 Pa) matrices due to the fact that cells stayed in the G1 phase cell cycle phase longer (Tilghman *et al*., 2012). Indeed, their results using cell lysates with ATPlite assay and protein synthesis assays confirmed their hypothesis. The substrates used in those experiments were limited to polyacrylamide gels that have a large rigidity/flexibility range, but it is not physiological. In our approach, we used collagen monolayers prepared at two different concentrations of 1.2 mg/mL and 3.0 mg/mL Second harmonic generation (SHG) images were taken to measure the fiber thickness, and density was measured using image correlation spectroscopy (ICS). Previously in our lab, we have shown that mechanical properties of collagen obtained through SHG and ICS correlated to those obtained by rheology or scanning electron microscopy images (Chiu *et al*., 2013; Raub *et al*., 2008). For this analysis, the ω_o_ value gives the waist of the auto correlation function and based on the size of the point spread function of the laser (∼0.3 µm at the waist). A larger ω_o_ indicates thicker fibers. 1/G(0) quantified the density of the matrix which is the height of the auto correlation function extrapolated from the first measured point. A smaller 1/G(0) value corresponds to denser matrices. 3.0 mg/mL and 1.2 mg/mL collagen substrates showed similar average values of ω_o_ of 2.55 and 2.29 (Figure 1C). However, the 3.0 mg/mL collagen has a significantly larger average value of 1/G(0) of 5.54 compared to that of the 1.2 mg/mL collagen at 1.09 (Figure 1D). This confirms that the 3.0 mg/mL collagen substrates have a denser network of collagen although their fiber thicknesses are similar. Rheology measurements were also done to quantify the modulus of the substrates. 3.0 mg/mL and 1.2 mg/mL collagen substrates were measured and have averages of 38.12 Pa and 5.66 Pa, respectively (Figure 1B).

**Figure 1:**
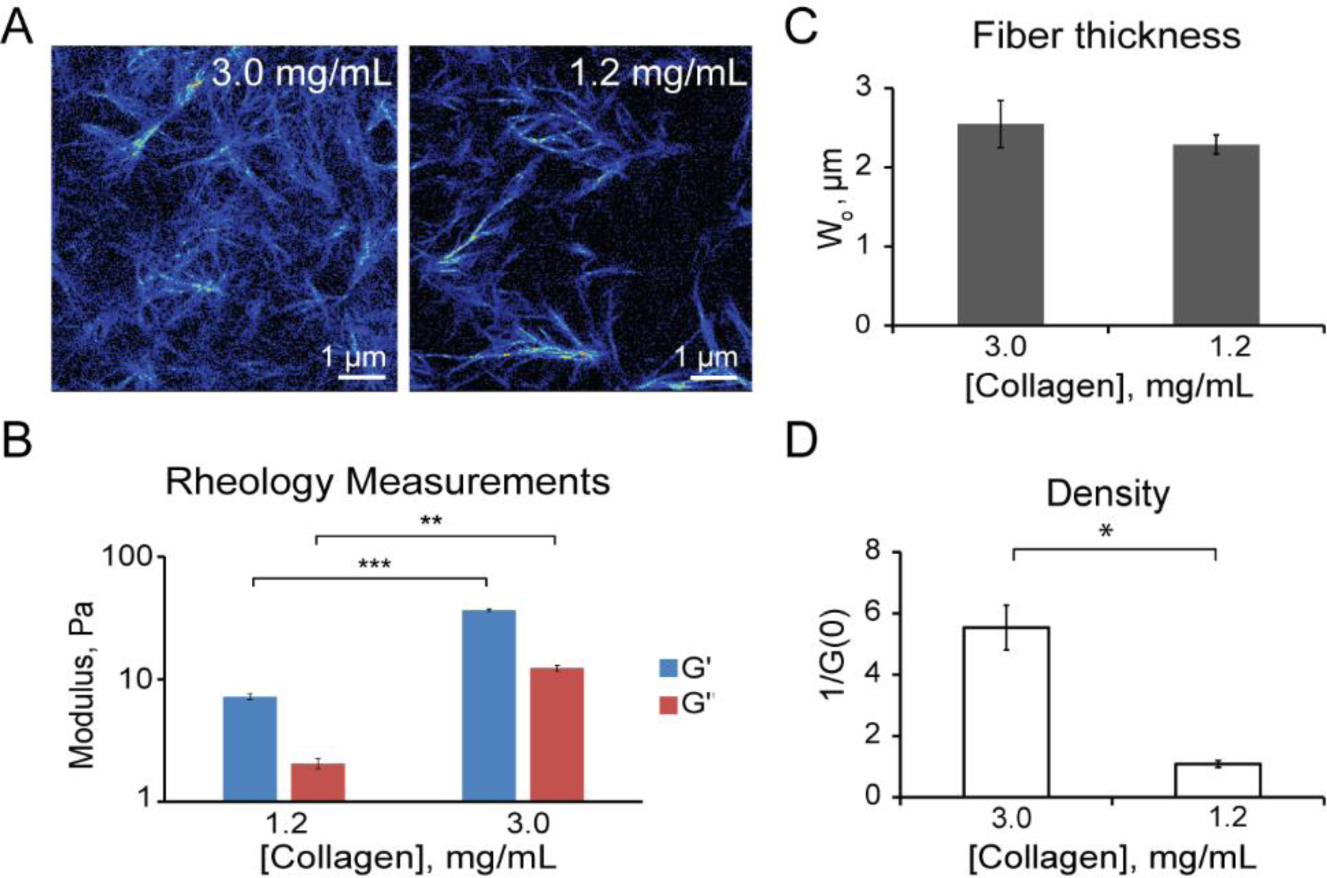
Quantification of collagen substrates. (A) Second harmonic generation images of 3.0 mg/mL and 1.2 mg/mL collagen substrates. (B) Modulus of collagen substrates at 10% strain and 1Hz (C) Quantification of fiber size (ω_o_) and (D) density of collagen substrates. * p<1e-3, ** p<1e-26, *** p<1e-57.

### MDA-MB231 cells shifts towards glycolytic signatures on stiffer collagen substrates

By measuring NADH fluorescent lifetimes with FLIM, we were able to non-invasively determine spatial shifts in metabolism of different cell lines in response to collagen substrate stiffness. NADH has two different lifetimes when it is free in the cytosol, ∼0.4 ns, or bound to a protein, ranging from ∼1.4 ns to 9 ns (Jameson *et al*., 1989; Lakowicz *et al*., 1992; Skala *et al*., 2007). Thus, we are able to distinguish the ratio of free and bound NADH at each pixel. For our studies, the lifetime of NADH bound to lactate dehydrogenase (LDH, ∼3.4 ns) is used when quantifying the population of bound NADH, although there are many other possible enzymes (Datta *et al*., 2015; Ma *et al*., 2016). The lifetime decay measured is Fourier transformed and displays a graphical representation of fluorescence lifetime on the phasor plot where all single exponential decay lifetimes are plotted on the semi-circle (called the universal circle) and all multi-exponential lifetimes are inside the semicircle representing the sum of linear combinations of single-exponential lifetimes. Figure 2A depicts the fluorescence lifetime (0.4 ns) of free NADH and 100% bound (3.4 ns) NADH to LDH. If the binding is not complete, we can calculate the ratio of bound to free NADH from the pure free and bound linear combination of lifetimes. In our control, the population of percentage bound to NADH was 75%. This linear trajectory is indicative of GLY and OXPHOS state in live cells (Digman *et al*., 2008a; Stringari *et al*., 2012).

**Figure 2:**
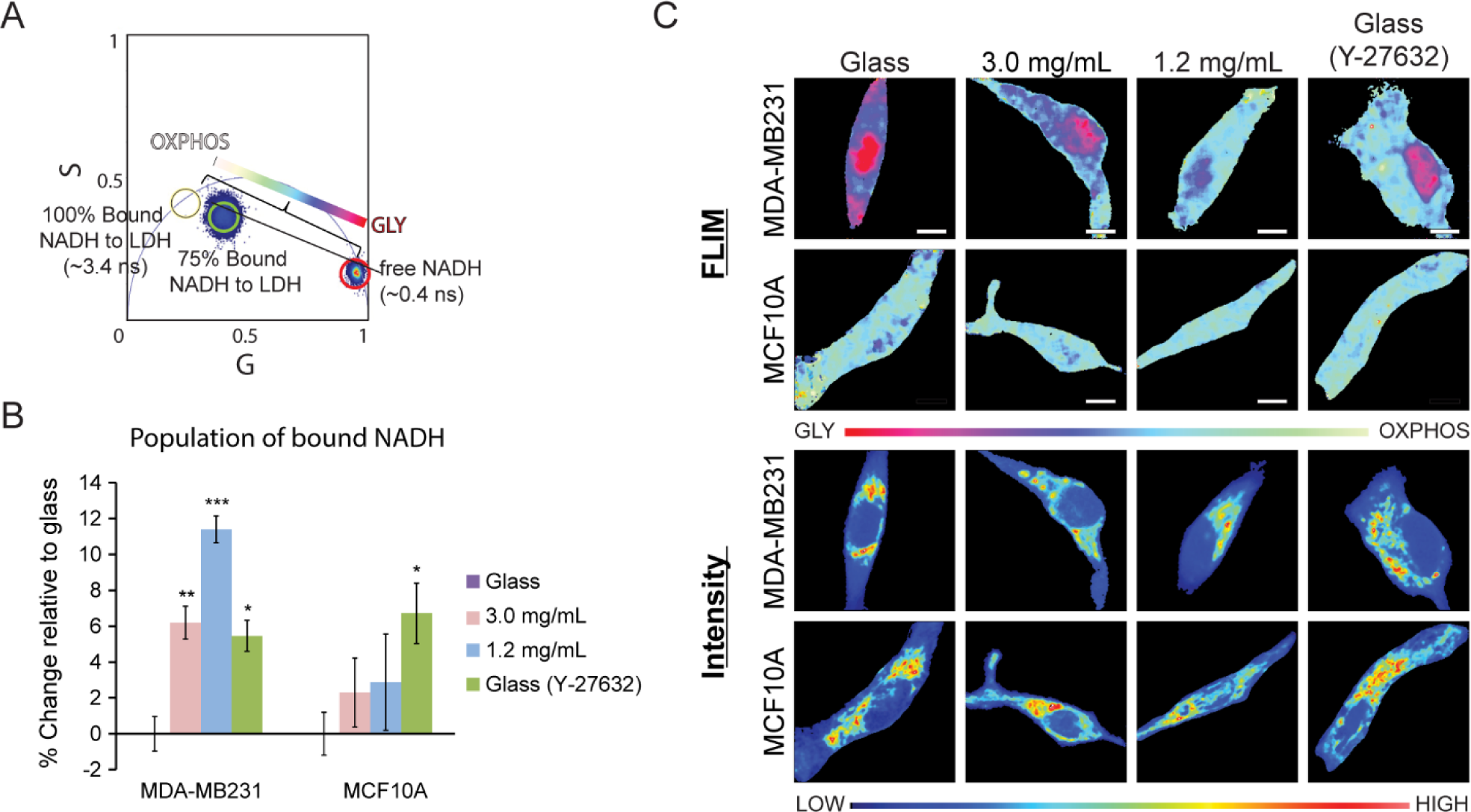
Metabolic indexes of MDA-MB231 and MCF10A cells on various substrate stiffnesses. (A) An increased population of bound NADH to LDH (long lifetime NADH, cyan) is indicative of a more OXPHOS signature while an increased population of free NADH (short lifetime NADH, red) would indicate GLY. These two extremes create a linear “M-trajectory” where mixed population of bound and free NADH, for example 75%, will lie between these two points. (B) Percent increase of bound NADH in MDA-MB231 and MCF10A cells relative to glass. (C) Colored images of FLIM of NADH and the average intensity of NADH within MDA-MB231 and MCF10A cells. *p<0.05, **p<0.01, and ***p<1e-5 by Student’s t-test. Scale bar: 5 µm.

Highly invasive MDA-MB231 cells were seeded on collagen and glass substrates to observe changes in free:bound ratios of NADH. In addition, the free:bound ratio of NADH in non-tumorigenic breast cells MCF10A were used as a control for a non-tumorigenic cell line. MDA-MB231 cells showed a 6.2% and 11.4% increase of bound NADH on 3.0 mg/mL and 1.2 mg/mL substrates, respectively, relative to those on glass (Figure 2B). This indicates that as substrate stiffness increased, the MDA-MB231 cells shifted from OXPHOS (white/cyan) to GLY (pink/red) (Figure 2C). MDA-MB231 cells on glass were then treated with 10 µM Y-27632, a ROCK inhibitor that decreases cell contractility, to assess if inhibiting the cell’s mechanosensing ability would shift the metabolism towards OXPHOS. Indeed, we see a significant change in the metabolic index (5.7% increase in bound NADH) in these cells when Y-27632 was added (Figure 2B). Similar results were seen on MDA-MB231 cells on glass when treated with 3.5 µM blebbistatin, showing a 14.4% increase relative to untreated glass samples (Supplementary Figure S1). We also used both treatments on MDA-MB231 cells plated on 1.2 mg/mL and 3.0 mg/mL collagen and detected an increase in the bound NADH population as well (Supplementary Figure S2). MCF-10A cells did not show this shift in metabolism in response to substrate stiffness, but did show an increase in the population of bound NADH with the addition of Y-27632 on glass (Figure 2B&C). We also examined other cancer cell lines to determine if they also had the same response to collagen stiffness. These results are shown below.

Melanoma, A375MM, and glioblastoma, U251MG, cell lines showed different results than that of the MDA-MB231 cells. A375MM cells shows no significant change in free:bound ratio of NADH when on the 1.2 mg/mL or 3.0 mg/mL collagen substrates or glass (Figure 3A&B). We noticed that these cells did not adhere as well on the collagen substrates due to their round morphology (Figure 3C) which could be the reason why there was no change in metabolism as seen in the MDA-MB231 cells. This further supports our hypothesis that the mechanosensing pathway plays an important role in cancer cell metabolism. On the other hand, the U251MG cells showed opposite changes to that of MDA-MB231 cells with 3.4% and 1.6% decrease in bound NADH relative to glass on 3.0 mg/mL and 1.2 mg/mL collagen, respectively (Figure 3B). Thus, they showed increased glycolytic signatures as substrate stiffness decreased, although the changes were not significant. Previous studies have also shown that MDA-MB231 and U251MG have opposite trends of matrix degradation and invadopodia formation when they are cultured in different media supplemented with GLY or OXPHOS inhibitors (Van Horssen *et al*., 2013). This may explain our results, but further studies will need to be conducted to confirm them.

**Figure 3:**
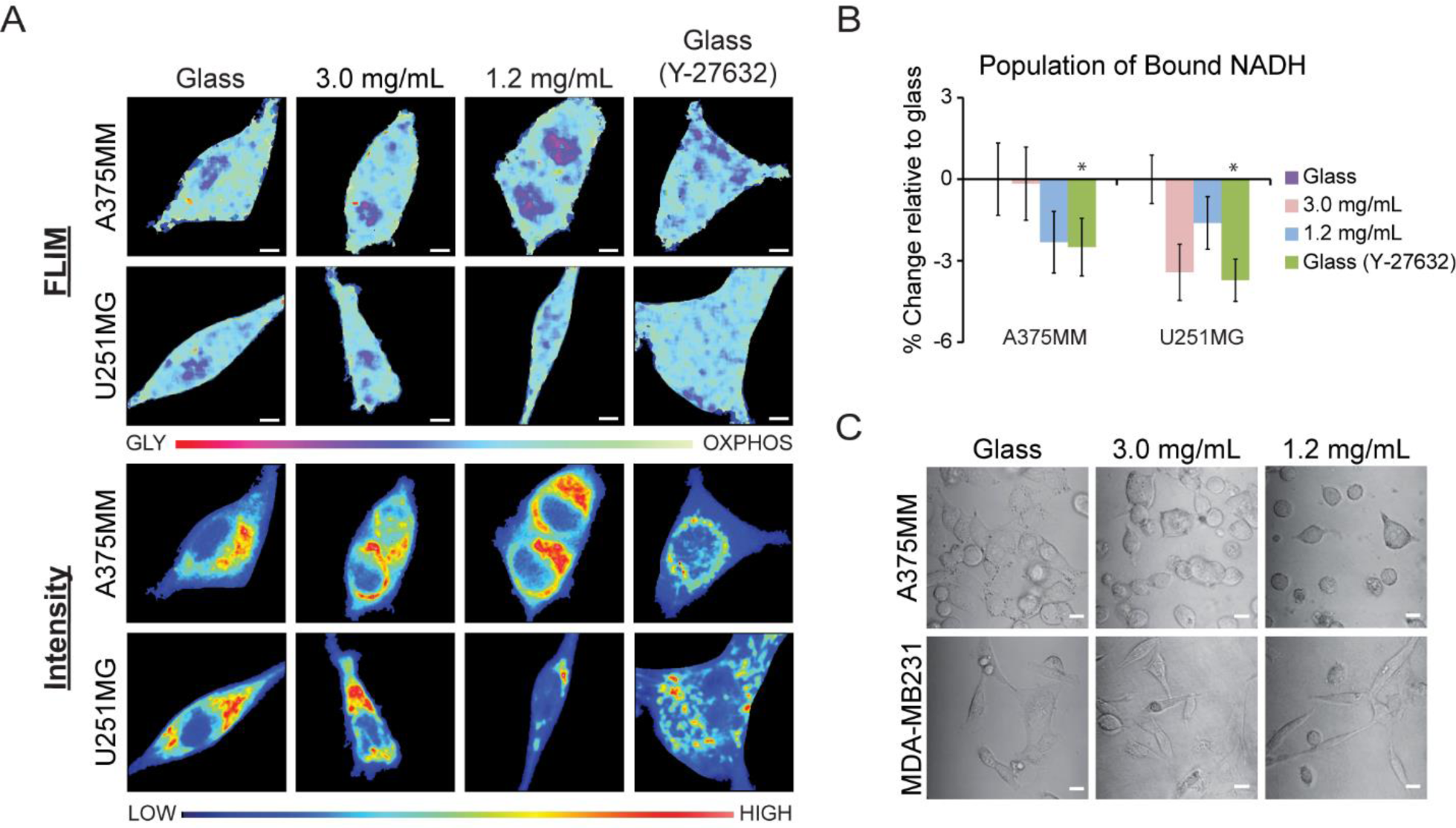
Metabolic indexes of A375MM and U251MG cells on various substrate stiffnesses. (A) FLIM and average intensity images of NADH within A375MM melanoma and U251MG glioma cell lines. (B) quantification of the percent change of NADH within A375MM and U251MG cell lines on various substrates relative to those on glass. (C) Transmitted optical images of A375MM and MDA-MB231 cells on different substrates. *p<0.05, by Student’s t-test. Scale bar: 5 µm.

We isolated the metabolic phasor signature of the nucleus and the cytoplasm and compare them here across all surfaces in each cell line (Supplementray Figure 4). Generally, the nucleus of the cell lines has a more GLY signature than the cytoplasm, but this did not significant effect the results we found when looking at the entire cell within MDA-MB231 and MCF10A cells. However, within A375MM cells seeded on 3.0 mg/mL or 1.2 mg/mL collagen substrates, their populations of bound NADH were similar; but when looking at their nuclei and cytoplasmic, we began to see a separation between the two conditions, especially in the nuclei alone. The nuclei in A375MM cells on 3.0 mg/mL collagen substrates show a shift towards GLY where their population of bound NADH decreased by 24.5% compared to those on glass. U251MG cell nuclei metabolic indices were similar on all surfaces except for those on glass and treated with blebbistatin, which show a 10.6 decrease in the population of bound NADH.

**Figure 4:**
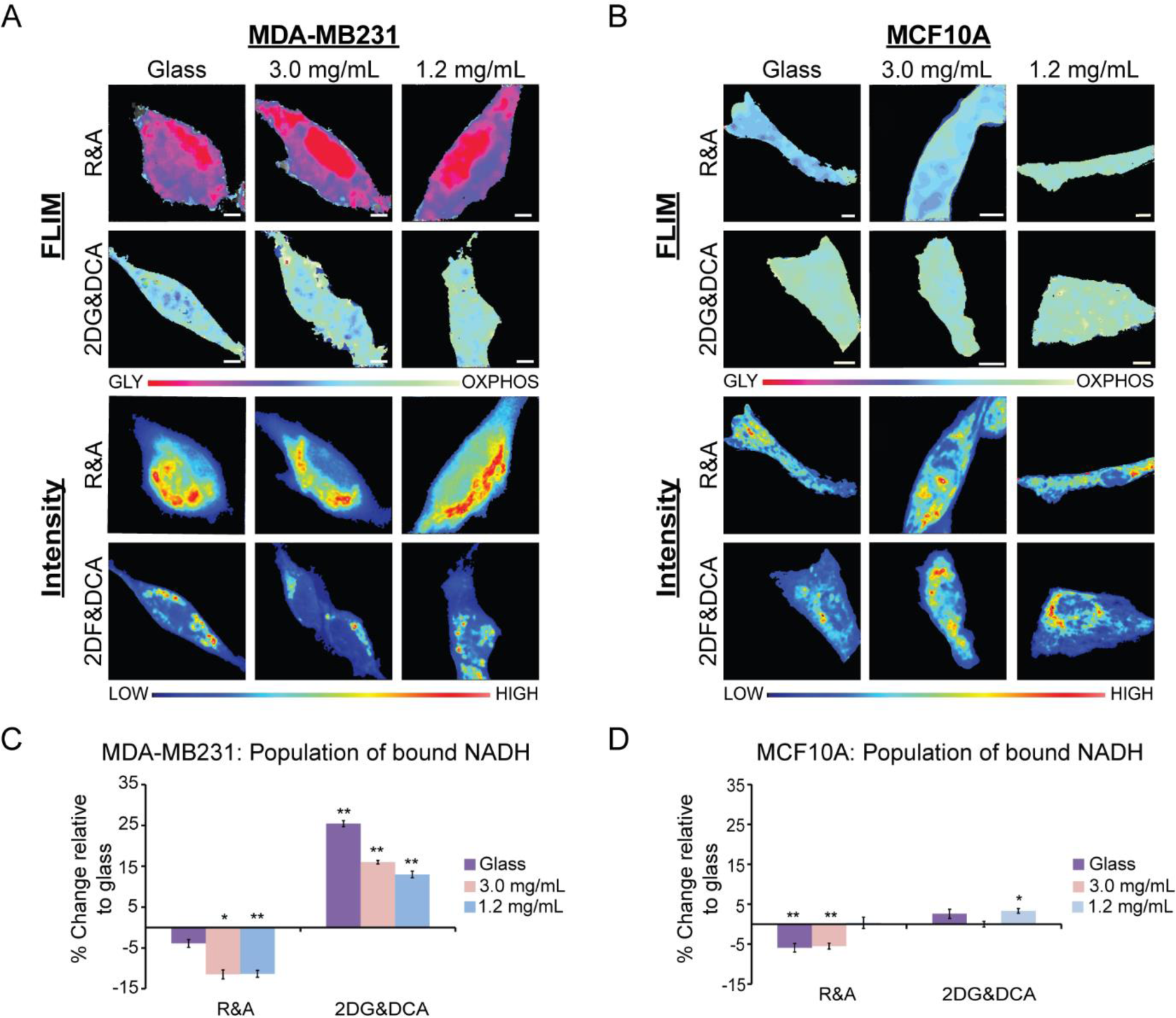
MDA-MB231 and MCF10A metabolic indexes when treated with metabolic inhibitors. (A) MDA-MB231 and (B) MCF10A cells treated with rotenone and antimycin A (R&A) or 2-deoxy glucose and dichloroacetate (2DG&DCA) for OXPHOS or GLY inhibition, respectively. FLIM images show the shifts of metabolic indexes with respective intensity images below. (C) Quantification of the change of bound NADH in MDA-MB231 and (D) MCF10A cells relative to untreated cells. *p<1e-3, **p<1e-4 or less by Student’s t-test. Scale bar: 5 µm.

The rest of this report will focus mainly on the MDA-MB231 and MCF10A cells to compare results of cancerous and non-tumorigenic cell lines. In order to confirm that the fraction of free:bound ratio of NADH is modulated through substrate stiffness, we conducted metabolism inhibition studies of each cell line when seeded on both collagen and glass substrates.

### Metabolism inhibition studies confirm that GLY and OXPHOS are modulated by substrate stiffness

In order to ensure that the collagen density alters metabolism in MDA-MB231 and MCF10A cells and that these changes correlate with lifetime positions along the M trajectory, we treated these cell lines with oxidative phosphorylation and glycolysis inhibitors. We treated MDA-MB231 cells with 50 mM 2-deoxyglucose and 100 mM dichloroacetate (2DG&DCA) for GLY inhibition and 50 nM rotenone and 50 nM antimycin A (R&A) for OXPHOS. 2DG&DCA treatment showed an increased population of bound NADH of 25.43%, 16.01% and 12.98% for cells plated on glass, 3.0 mg/mL collagen, and 1.2 mg/mL collagen surfaces, respectively, relative to untreated cells (Figure 4C). When MDA-MB231 cells were treated with R&A on 3.0 mg/mL and 1.2 mg/mL collagen substrates, they showed significant decreases in the population of bound NADH, p<0.05, of 11.5% and 11.35%, respectively, relative to untreated cells. However, there was no change when these cells plated on glass were inhibited with R&A. These treatments were also applied to MCF10A cells where we observed a significant decrease in the population of bound NADH only on 3.0 mg/mL collagen substrates, 5.52%, and glass, 5.93%, when treated with R&A (Figure 4D). No changes were seen on the 1.2 mg/mL collagen substrates. However, there was a significant increase of 3.32% on these cells when plated on 1.2 mg/mL collagen and treated with 2DG and DCA.

### Diffusion of focal adhesion proteins correlate to substrate stiffness

In order to correlate our finding of substrate stiffness to the stability of focal adhesions dynamics in cells, we analyzed the diffusion of talin-GFP, a protein associated with integrin mediated adhesions and the actin cytoskeleton, by using RICS, raster image correlation spectroscopy (Digman and Gratton, 2009; Digman *et al*., 2008b, 2009; Liang *et al*., 2017). In our previous work we demonstrated that increased spatio-temporal focal adhesion diffusion of proteins at the focal adhesions indicated active adhesion association and dissociation whereas stable adhesions had a steady state population. By measuring the diffusion of talin at the FA, we can observe the motility and stability of the adhesions. We hypothesized that MDA-MB231 on stiffer matrices would show a decreased diffusion of talin-GFP due to increased FA stability. We imaged MBA-MB231 cells transiently transfected with talin-GFP and observed an increase of diffusion with substrates stiffness (Supplementary Figure S5b). Talin-GFP of MDA-MB231 cells on glass had a diffusion rate of 0.71 µm^2^/sec where the cells on the 3.0 and 1.2 mg/mL collagen substrates had rates of 3.40 µm^2^/sec and 4.62 µm^2^/sec, respectively. Cells were treated with Y-27632, a specific inhibitor of kinase (ROCK) and cell contractility, to the disruption of the mechanical tension (Martin *et al*., 2016). These cells had an increased diffusion of talin-GFP of 2.04 µm^2^/sec.

## Discussion

The Warburg effect is the hallmark for cancer cell metabolism, described as an oncogene-directed glycolytic state even when oxygen is present (Vander Heiden *et al*., 2009; Liberti and Locasale, 2016; Ward and Thompson, 2012). This could be due to the high turnover of ATP production through glycolysis for energy production with glucose, although the alternative process of oxidative phosphorylation creates more product of energy per glucose molecule. Cancer cell invasion has also been shown to be modulated by changes in metabolism through changing cell media additives for energy consumption (Van Horssen *et al*., 2013; Scott *et al*., 2012) or the ECM stiffness (Artym *et al*., 2015; Gould and Courtneidge, 2014; Paszek *et al*., 2005; Provenzano *et al.*, 2008; Seewaldt, 2014). However, the link between ECM stiffness and cell metabolic state (OXPHOS or GLY) is not clear. Recent studies that look at alterations of metabolism use invasive biochemical assays that do not report the spatial heterogeneous changes of metabolic response within cellular compartments, nor their cellular metabolic state. Our study used FLIM of NADH to measure real-time metabolic indexes of different cancer cell types in response to collagen substrate stiffness. This allows us to characterize which metabolic process occurs with pixel resolution in live cells.

We have found that MDA-MB231 cells on substrates on two different collagen stiffnesses have an increasing population of free NADH, showing a more glycolytic signature of metabolism. This correlates with previous studies in that breast cancer cells have increased migration and aggressiveness within stiffer collagen matrices (Haage and Schneider, 2014; Levental *et al*., 2009; Paszek *et al*., 2005). In addition, cancer cells undergo aerobic glycolysis for energy production, thus increased substrate stiffness could be a contributor to the metabolic shift towards glycolysis (Liberti and Locasale, 2016; Morris *et al*., 2016). The ECM plays a key role in the cancer cell’s mechanosensing pathway through integrin signaling, and there is increasing evidence that this regulates cell migration and matrix degradation (Alexander *et al*., 2008; Beaty *et al*., 2013; Chiu *et al*., 2013; Van Horssen *et al*., 2013). Increased ECM stiffness signals actin polymerization, increased integrin signaling, and stabilization of the focal adhesion complexes (Ciobanasu *et al*., 2013; Gardel *et al*., 2010; Hirata *et al*., 2014). This increase of integrin signaling has been shown to also upregulate the PI3K/AKT/mTOR pathways and possible metabolism switching in cancer cells (Ata and Antonescu, 2017; Caino *et al*., 2015; Lien *et al*., 2016; Yang *et al*., 2015). In addition, stimulating the actin-contractility of cells through external forces by shear flow or pulling of the cell membrane increases glucose uptake (Bays *et al*., 2017; Hayashi *et al*., 1998). Bays *et al.* has shown that this increased glucose uptake was also shown to increase ATP production for actin polymerization and stabilize E-cadherin contacts. Our results expanded on these studies by looking at the specific metabolic indexes of cancer cells when introduced to various substrate stiffness. We speculate that these same pathways are being activated in the MDA-MB231 cells and are stimulated passively through focal adhesion-mediated interactions with the ECM.

When cells grown on glass substrates were treated with Y-27632 or blebbistatin, we inhibited their ability to undergo contraction through myosin-II and caused focal adhesion detachment from the substrate (Martin *et al*., 2016). This in turn showed shifts in metabolic indexes from GLY to OXPHOS in MDA-MB231 and MCF10A cells. This confirms that it is through actin-mediated cell contractility that modulated these shifts in metabolism. Interestingly, the MDA-MB231 cells treated with Y-27632 had a similar NADH free: bound ratio as those grown on 3.0 mg/mL, which could mean that their degrees of contractility were similar to each the respective conditions. Those treated with blebbistatin had a much larger shift towards OXPHOS, surpassing the population of bound NADH of cells grown on 1.2 mg/mL collagen. Since blebbistatin directly affects myosin-II and is more potent than Y-27632, this was as expected. We also used RICS to measure the dynamics of the talin within MDA-MB231 cells to correlate these movements to adhesion stability. Previous studies have shown that stable adhesions have a developed mechanosensing network of actin, and these adhesions can grow in response to forces or increasing substrates stiffness (Bieling *et al*., 2016; Kim and Wirtz, 2013; Petit and Thiery, 2000; Wells, 2008). Thus, cells on the stiffest substrates should have the most stable adhesions and a decreased diffusion rate due to lack of unbinding. Increased FAK promotes glucose consumption and play a key role in the OXPHOS and GLY balance within cancer cells (Palorini *et al*., 2013). MDA-MB231 cells expressing talin-GFP were imaged over time on each of the substrates, and an area containing the focal adhesions was analyzed with RICS to extract the diffusion of those proteins. Our results show that the cells on the glass substrates contained the slowest diffusion rates of 0.71 µm^2^/sec and the fastest of 4.62 µm^2^/sec for cells grown on 1.2 mg/mL substrate. This supports our hypothesis that our softest substrates would give rise to faster moving adhesions due to their instability and thus constant unbinding and binding. In addition, it further supports that an upregulation of adhesion through substrates stiffness shifts MDA-MB231 cells towards a glycolytic signature. We saw that when these cells on the glass substrate were treated with Y-27632, had an increased diffusion of 2.05 µm^2^/sec from 0.71 µm^2^/sec.

The human melanoma cell lines, A375MM, in this study attached to 1.2 mg/mL and 3.0 mg/mL collagen substrates but did not spread as well as all other cells. This was an indication that their adhesions are not stable or favorable on these substrates (Calderwood *et al*., 2013; Cavalcanti-Adam *et al*., 2007; Massia and Hubbell, 1991) and their mechanosensing ability could have been compromised and reduced mitochondria activity (Ochsner *et al*., 2010). Consequently, this would fail to change the metabolic indexes of these cells as shown in our results; where the NADH free: bound of A375MM on 1.2 mg/mL or 3.0 mg/mL have no significant difference. This phenotype showed that there is a confirmed link between focal adhesion-mediated mechanosensing and cellular metabolism

We also measured the NADH free:bound ratio in glioblastoma cells, U251MG, to see if these trends existed in other cancer cell lines. It was found that they had an opposite trend than that of the MDA-MB231 cells. As substrate stiffness increased, the U251MG cells had a more OXPHOS metabolic index, or an increase in bound NADH. Studies done by Van Horssen *et. al* also showed that MDA-MB231 and U251MG cells had opposite trends in matrix degradation and invadopodia formation when their metabolism is altered either by galactose containing media to inhibit GLY or addition of oligomycin to inhibit OXPHOS (Van Horssen *et al*., 2013). It is important to note that there are many other possible uses for pyruvate aside from OXPHOS in the mitochondria. Downstream of GLY are intermediates of the tricarboxyl acid cycle, such as citrate, which allows for synthesis of lipids, proteins and nucleic acids, a demand for highly proliferating cells (DeBerardinis *et al*., 2008). There is also fatty acid synthesis through citrate which is shown to correlate with the formation of invadopodia, which are actin rich protrusions used for matrix invasion (Gould and Courtneidge, 2014; Morris *et al*., 2016; Scott *et al*., 2012). In addition, the pentose phosphate pathway could further increase the GLY and fraction of free NADH. All of the pathways mentioned could be elevated within MDA-MB231 cells, which could contribute to their difference in metabolic trends from the U251MG cells when ECM stiffness is varied.

The MB231 cells have a significant decrease in the fraction of bound NADH when plated on glass, 3.0 mg/mL and 1.2 mg/mL collagen, respectively. We confirmed that the changes with metabolic trajectory were reflective in cellular metabolism using the OXPHOS and GLY inhibitors. When these inhibitors were added, cells shifted their metabolism accordingly to their inhibitors but there were no significant metabolic differences across substrate density within these changes (Supplementary Figure S3A). However, the MCF10A cell lines did not show any significant changes across substrate densities in the untreated conditions. They did show substrate sensitivity only when OXPHOS was inhibited. When R&A was added to inhibit OXPHOS to the 3.0 mg/mL and glass substrates in MCF10A cells, there was a maximum decrease to around 64% of the population of bound NADH; however, those on 1.2 mg/mL collagen showed no significant change (Supplementary Figure S3B). This could mean that on stiffer substrates, these cells were more susceptible to metabolic changes when introduced to inhibitors. Additionally, this could also indicate that the metabolism of the MCF10A cells was behaving more like the MDA-MB231 cells on stiffer matrices. When 2DG&DCA was added to inhibit GLY in MCF10A cells, we see a significant increase of the population of bound NADH to around 71.2% when grown on 1.2 mg/mL collagen substrate. Since OXPHOS and softer substrates is preferable for the MCF10A cells, this could mean that this ECM provides an extra boost towards OXPHOS pathway when GLY is inhibited.

The phasor approach to FLIM of NADH allows isolation of the metabolic signature within sub-cellular compartments of the cells. Here, we focused on comparing the nuclei and cytoplasm of MDA-MB231, MCF10A, A375MM and U251MG cell lines (Supplementary Figure S4). We were able to see that the metabolic shifts within the nuclei and cytoplasm of MDA-MB231 and MCF10A cells are similar to their whole cell signature. However, within A375MM cells we were able to make distinctions of the population of bound NADH between surfaces, which were not detected when averaging over the entire cell. The nuclei of A375MM cells on 3.0 mg/mL collagen substrates has a significant decrease in the population of bound NADH with respect to those on glass. Thus, looking at the nuclear metabolic indexes can separate subtle changes that are hidden in whole cell readings. These distinctions seen could be due to nuclear processes, such as transcription or DNA repair, which has also been shown to affect the ratio of bound and free NADH (Aguilar-Arnal *et al*., 2016; Wright *et al*., 2012).

## Conclusion

We have shown that focal adhesion-mediated contractility modulates cell metabolism in MDA-MB231 cancer cells. With the use of FLIM of NADH, we were able to measure metabolic changes of cancer cell lines MDA-MB231, A375M and U251MG and within non-tumorigenic line MCF10A. Particularly in breast cancer MDA-MB231 cell lines, we saw that stiffer substrates shifted cells to have a more glycolytic metabolic signature due to their increased population of free NADH. However, in non-tumorigenic breast cells MCF10A, we did not see any changes in metabolism across all substrates. Further studies by inhibiting myosin-II contractility increased the population of bound NADH MDA-MB231 cells across all surfaces and confirmed our hypothesis. In addition, an increase in substrate stiffness was seen to results in less dynamic talin proteins and more stable adhesions, indicating an established mechanosensing network. This further supports our assertion that ECM mediated adhesions are upregulated due to substrate stiffness and modulates metabolic signatures.

We also confirmed that the changes in NADH free:bound ratio in MDA-MB231 and MCF10A cells were due to GLY or OXPHOS by inhibiting these pathways with dichloroacetate and 2-deoxyglucose or rotenone and antimycin A, respectively. With our results combined with what is known in literature, there is a relationship between the mechanosensing and metabolism pathway in cancer cells, and both play a critical role in regulating cancer invasiveness. This provides insight to develop therapies which target mechanosensing abilities of cancer cells to revert their metabolism similar to a more non-tumorigenic cell type or decrease their invasiveness.

## Materials and Methods

### Cell culturing and transfections

MDA-MB231 and A375MM cells were cultured in Dulbecco’s Modified Eagle’s Medium (DMEM) with high glucose, L-glutamate, and sodium pyruvate (Genesee Scientific, San Diego, CA) supplemented with 10% heat inactivated Fetal Bovine Serum (Thermofisher Scientific, USA) and 1% Penicillin-Streptomycin 100X Solution (Genesee Scientific, San Diego, CA). MCF10A cells were cultured in DMEM/F12 with high glucose, sodium pyruvate and L-glutamine (Thermofisher Scientific, USA) supplemented with 5% horse serum (Thermofisher Scientific, USA), 20 ng/mL epidermal growth factor, 0.5 mg/mL Hydrocortisone (Sigma-Aldrich, St. Louis, MO), 100 ng/mL cholera toxin (Sigma-Aldrich, St. Louis, MO), 10 µg/mL insulin (Sigma-Aldrich, St. Louis, MO), and 1% Penicillin-Streptomycin 100X Solution (Genesee Scientific, San Diego, CA). U251MG cells were cultured in DMEM/F12 with high glucose, sodium pyruvate and L-glutamine (Thermofisher Scientific, USA) supplemented with 10% heat inactivated Fetal Bovine Serum (Thermofisher Scientific, USA) and 1% Penicillin-Streptomycin 100X Solution (Genesee Scientific, San Diego, CA). All cell lines were incubated at 37^°^C, 5% CO_2_.

When cell lines which required transfections of talin-GFP (Addgene, Cambridge, MA) were seeded in a 6-well plate at 0.25−10^−6^ cells/well overnight at 37^°^C, 5% CO_2_. Lipofectamine 3000 (Thermofisher Scientific, USA) was used for transfections following manufacture protocol. Briefly, 100 µL of Opti-mem (Thermofisher Scientific, USA) and 5µL of Lipofectamine 3000 were mixed in a microcentrifuge tube. In a second microcentrifuge tube, 100 µL of Opti-mem, 1 µg of DNA plasmid, and 2 µL/µg of P3000 reagent were mixed together. Both tubes were allowed to sit for 5 minutes at room temperature and then combined. After the transfection mixture was allowed to sit for 25 minutes at room temperature, it was added dropwise to the cells.

### Collagen substrate monolayers

Substrates were made on 35 mm glass bottom imaging dishes which were treated for 5 minutes with UV-ozone. 1% v/v of 3-aminopropyltriethoxysilane (APTES) in deionized water were added and allowed to sit for 25 minutes at room temperature. The dishes were washed thoroughly with deionized water and brought to the biosafety hood for sterile handling. 1 mL of sterilized MilliQ water used to rinse the dish before collagen was added.

For collagen preparation, microcentrifuge tubes and reagents used were kept on ice for as long as possible while handling. Collagen I from rat tail (Corning, Corning, NY) was diluted with deionized water such that the final concentration was either 1.2 mg/mL or 3.0 mg/mL. 100 µL of 10X phosphate buffer saline (Thermofisher Scientific, USA) containing phenol red was added dropwise while vortexing as a pH indicator. 5N NaOH was then added dropwise to the mixture with periodic vortexing until the solution became a slight pink (pH∼7). The collagen was then added to the treated imaging dish (total volume of 1 mL) and incubated at 20^°^C for 1 hour and then at 37^°^C, 5% CO_2_ overnight. 0.25−10^−6^ cells were then added to each dish the next day and then allowed to incubate at 37^°^C, 5% CO_2_ overnight again before imaging took place.

### Inhibition studies

Contractility inhibition was done with Y-27632 (Selleckchem, US) or blebbistatin (Sigma-Aldrich, St. Louis, MO) at a working concentration of 10 µM and 3.5 µM, respectively. Each inhibitor was incubated for 10 minutes at room temperature before conducting FLIM/NADH imaging. MDA-MB231 or MCF10A cells were treated with sodium dichloroacetate (Sigma-Aldrich, St. Louis, MO) and 2-deoxyglucose (Sigma-Aldrich, St. Louis, MO) at a working concentration of 100 mM and 50 mM, respectively, for 6 hours at 37°C, 5% CO_2_ to inhibit glycolysis (Cinco *et al*., 2016). Similarly, cells were treated for 10 minutes with 50 nM rotenone and 50 nM antimycin A at 37°C, for oxidative phosphorylation inhibition studies before NADH lifetimes were measured.

### Characterization of collagen substrates

Rheology measurements of collagen substrates were conducted to obtain the storage (G’) and loss (G”) moduli. The collagen substrates were pre-made on 15 mm glass slides that were treated with UVO and APTES as described above. Collagen solutions of 1.2 mg/mL or 3.0 mg/mL was carefully pipetted onto the round glass slides and allowed to incubate at 20^°^C for 1 hour and then at 37^°^C overnight for 2 nights to mimic culturing conditions for imaging. The glass sides were placed on the stage of the AR-G2 rheometer (TA Instruments, New Castle, DE) that was kept at a constant temperature of 37°C. A sand-blasted parallel plate geometry with a diameter of 25 mm was lowered to a gap distance of 0.3 mm so that it was in close contact with the surface of the collagen. Rheology measurements were conducted at a constant sinusoidal frequency of 1 Hz and 10% peak-to-peak strain and outputs of G’ and G” were recorded for a total of 10 minutes. A data point was taken every 60 seconds.

Second harmonic generation imaging of collagen substrates were conducted to characterize substrate density as previously described (Chiu *et al*., 2013; Raub *et al*., 2007). Briefly, 2-photon excitation at 900 nm was used to generate second harmonics of collagen and collected with a bandpass filter at 460/80 nm with external photon-multiplier tubes (H7422P-40, Hamamatsu, Japan) and FastFLIM FLIMBox (ISS, Champaign, IL).100 frames were collected and analyzed using image correlation spectroscopy on SimFCS (LFD, UCI). Spatial correlations were applied to each pixel at coordinate (*x, y*) of the complied SHG images with equation (1):

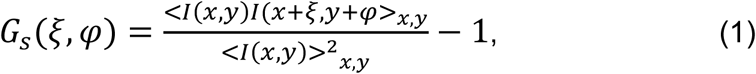

where *I* is the intensity. ξ and φ are the spatial shifts in the *x* and *y* directions, respectively.

### Confocal and fluorescent lifetime imaging acquisition and analysis

FLIM images for MDA-MB231, U251MG and A375MM cells were imaged on the Zeiss LSM 710 (Carl Zeiss, Jena, Germany), LSM 880 (Carl Zeiss, Jena, Germany), and Olympus Fluoview respectively. MCF10A cells were also imaged on the LSM710. Metabolism inhibition studies of MDA-MB231 and MCF10A cells were imaged on the Olympus Fluoview. Images (256−256 pixel size) were taken at a pixel dwell time of 25.21µs, 16.38 µs, and 20 µs for the LSM710, LSM880, and Fluoview, respectively. All microscope systems were coupled to with a two-photon Ti: Sapphire laser (Spectra-Physics MaiTai, Mountain View, CA) for NADH excitation at 740 nm with an Olympus 40X/0.8 NA water objective. The emission was separated at 690 nm in all systems followed by two bandpass filter at 460/80 nm and 540/50 nm with a with a dichroic mirror 495 nm long-pass filter. The signal was collected with an external photomultiplier tube (H7422P-40, Hamamatsu, Japan). A320 FastFLIM FLIMbox (ISS, Champaign, IL) was used to acquire the frequency domain of the lifetime of NADH until enough statistics was obtained. Images of coumarin-6 in ethanol were also taken as reference and calibration for FLIM measures across all microscopes.

SimFCS (LFD, UCI) was used to analyze the fluorescent lifetime of NADH at every pixel. The lifetime decay at each pixel was Fourier transformed and plotted on a phasor plot as previous described where each point on the phasor represents one pixel (Digman *et al*., 2008a). Each cell’s cluster of phasor points were averaged to obtain their S, G, and fraction bound value. The fraction of bound NADH is calculated based on the fact that any two points on the phasor plot (e.g. 100% free NADH and 100% bound NADH to LDH) can be connected by a line and any points along that line will be a linear representation of the two extremes. Thus, the experimental data will exist between 100% free NADH and 100% bound NADH, signifying a samples that has a mixture of free and bound NADH. Those points that are closer to the phasor of bound NADH to LDH will have a higher population of bound NADH.

### Focal adhesion imaging analysis

MBA-MB231 cells were transfected with talin-GFP and imaged on the Olympus FV1000 or Zeiss 880 microscope. 100 frames were taken at a 256−256 pixels where the pixel size was no more than 90 nm with a dwell time of 4.10 us/pixel for the Olympus or LSM 880, respectively. These images were analyzed with SimFCS using the raster image correlation spectroscopy to a region of interest (ROI) of 64−64 pixels of where the focal adhesions were located. The RICS routine as previously described (Digman and Gratton, 2009; Digman *et al*., 2008b; Rossow *et al*., 2010). A moving average subtraction of 10 frames was applied to analysis to account for any bulk cell motion. ω_o_, or the waist of the point spread function, was estimated as 0.3 µm. After generation of the auto correlation function, a fit was applied with the correct imaging parameters and the diffusion of the talin-GFP molecules within the ROI.

### Statistical analysis

Statistical significance was determined for experiments with the Student’s t-test (two-sample, unequal variance) in Microsoft Excel.

### Author Contributions

E.J.M. conducted all studies relating to cell mechanosensing and metabolism along with their FLIM analysis. Metabolism inhibition image collection and FLIM analysis were done by G.M. Focal adhesion protein, second harmonic generation of collagen, and rheology measurements and analysis were conducted by E.J.M. Collagen substrates were prepared by E.J.M. and G.M for their respected experiments. E.J.M., G.M., A.F.Y., and M.A.D. contributed to data interpretation and manuscript preparation.

## Acknowledgements

The authors sincerely thank Dr. Enrico Gratton, director of the Laboratory for Fluorescent Dynamics, for his advice on image correlation spectroscopy and raster image correlation spectroscopy on focal adhesions; Michael Murata, Andrew Trinh, Ning Ma, and Sara Sameni for their advice on NADH FLIM data; Jeremy Jacinto for organizing figures for this manuscript; the Daniela Bota Lab at UC Irvine for donating U251MG cells, and the Feng Liu-Smith lab at UC Irvine for donating A375MM cells. This work was supported by the National Institutes of Health grant P41-RRO3155 and in part by the American Cancer Society Institutional Research Grant 129801-IRG-16-187-13-IRG from the American Cancer Society.

## References

Aguilar-Arnal, L., Ranjit, S., Stringari, C., Orozco-Solis, R., Gratton, E., and Sassone-Corsi, P. (2016). Spatial dynamics of SIRT1 and the subnuclear distribution of NADH species. Proc. Natl. Acad. Sci. U. S. A. 113, 12715–12720.

Alexander, N.R., Branch, K.M., Parekh, A., Clark, E.S., Iwueke, I.C., Guelcher, S.A., and Weaver, A.M. (2008). Extracellular matrix rigidity promotes invadopodia activity. Curr. Biol. 18, 1295–1299.

Artym, V. V, Swatkoski, S., Matsumoto, K., Campbell, C.B., Petrie, R.J., Dimitriadis, E.K., Li, X., Mueller, S.C., Bugge, T.H., Gucek, M., et al. (2015). Dense fibrillar collagen is a potent inducer of invadopodia via a specific signaling network. J. Cell Biol. 208, 331–350.

Ata, R., and Antonescu, C.N. (2017). Integrins and Cell Metabolism: An Intimate Relationship Impacting Cancer. Int. J. Mol. Sci. 18.

Bays, J.L., Campbell, H.K., Heidema, C., Sebbagh, M., and DeMali, K.A. (2017). Linking E-cadherin mechanotransduction to cell metabolism through force-mediated activation of AMPK. Nat. Cell Biol. 19, 724–731.

Beaty, B.T., Sharma, V.P., Bravo-Cordero, J.J., Simpson, M.A., Eddy, R.J., Koleske, A.J., and Condeelis, J. (2013). β1 integrin regulates Arg to promote invadopodial maturation and matrix degradation. Mol. Biol. Cell 24, 1661–1675, S1-11.

Bieling, P., Li, T.-D., Weichsel, J., McGorty, R., Jreij, P., Huang, B., Fletcher, D.A., and Mullins, R.D. (2016). Force Feedback Controls Motor Activity and Mechanical Properties of Self-Assembling Branched Actin Networks. Cell 164, 115–127.

Bird, D.K., Yan, L., Vrotsos, K.M., Eliceiri, K.W., Vaughan, E.M., Keely, P.J., White, J.G., and Ramanujam, N. (2005). Metabolic mapping of MCF10A human breast cells via multiphoton fluorescence lifetime imaging of the coenzyme NADH. Cancer Res. 65, 8766–8773.

Bugyi, B., and Carlier, M.-F. (2010). Control of actin filament treadmilling in cell motility. Annu. Rev. Biophys. 39, 449–470.

Caino, M.C., Ghosh, J.C., Chae, Y.C., Vaira, V., Rivadeneira, D.B., Faversani, A., Rampini, P., Kossenkov, A. V, Aird, K.M., Zhang, R., et al. (2015). PI3K therapy reprograms mitochondrial trafficking to fuel tumor cell invasion. Proc. Natl. Acad. Sci. U. S. A. 112, 8638–8643.

Calderwood, D. a, Campbell, I.D., and Critchley, D.R. (2013). Talins and kindlins: partners in integrin-mediated adhesion. Nat. Rev. Mol. Cell Biol. 14, 503–517.

Cavalcanti-Adam, E.A., Volberg, T., Micoulet, A., Kessler, H., Geiger, B., and Spatz, J.P. (2007). Cell spreading and focal adhesion dynamics are regulated by spacing of integrin ligands. Biophys. J. 92, 2964–2974.

Chiu, C.-L., Digman, M.A., and Gratton, E. (2013). Cell matrix remodeling ability shown by image spatial correlation. J. Biophys. 2013, 532030.

Cinco, R., Digman, M.A., Gratton, E., and Luderer, U. (2016). Spatial Characterization of Bioenergetics and Metabolism of Primordial to Preovulatory Follicles in Whole Ex Vivo Murine Ovary. Biol. Reprod. 95, 129–129.

Ciobanasu, C., Faivre, B., and Le Clainche, C. (2013). Integrating actin dynamics, mechanotransduction and integrin activation: the multiple functions of actin binding proteins in focal adhesions. Eur. J. Cell Biol. 92, 339–348.

Cunniff, A.B., Mckenzie, A.J., Heintz, N.H., and Alan, K. (2016). AMPK activity regulates trafficking of mitochondria to the leading edge during cell migration and matrix invasion Department of Pathology Department of Pharmacology University of Vermont Cancer Center University of Vermont, Burlington, VT 05405, USA Co. Mol. Biol. Cell 27, 2662–2674.

Datta, R., Alfonso-García, A., Cinco, R., and Gratton, E. (2015). Fluorescence lifetime imaging of endogenous biomarker of oxidative stress. Sci. Rep. 5, 9848.

DeBerardinis, R.J., Lum, J.J., Hatzivassiliou, G., and Thompson, C.B. (2008). The Biology of Cancer: Metabolic Reprogramming Fuels Cell Growth and Proliferation. Cell Metab. 7, 11–20.

Desai, S.P., Bhatia, S.N., Toner, M., and Irimia, D. (2013). Mitochondrial localization and the persistent migration of epithelial cancer cells. Biophys. J. 104, 2077–2088.

Digman, M.A., and Gratton, E. (2009). Analysis of diffusion and binding in cells using the RICS approach. Microsc. Res. Tech. 72, 323–332.

Digman, M.A., Caiolfa, V.R., Zamai, M., and Gratton, E. (2008a). The phasor approach to fluorescence lifetime imaging analysis. Biophys. J. 94, L14–6.

Digman, M.A., Brown, C.M., Horwitz, A.R., Mantulin, W.W., and Gratton, E. (2008b). Paxillin dynamics measured during adhesion assembly and disassembly by correlation spectroscopy. Biophys. J. 94, 2819–2831.

Digman, M.A., Wiseman, P.W., Horwitz, A.R., and Gratton, E. (2009). Detecting Protein Complexes in Living Cells from Laser Scanning Confocal Image Sequences by the Cross Correlation Raster Image Spectroscopy Method. Biophys. J. 96, 707–716.

Estrella, V., Chen, T., Lloyd, M., Wojtkowiak, J., Cornnell, H.H., Ibrahim-Hashim, A., Bailey, K., Balagurunathan, Y., Rothberg, J.M., Sloane, B.F., et al. (2013). Acidity generated by the tumor microenvironment drives local invasion. Cancer Res. 73, 1524–1535.

Gardel, M.L., Schneider, I.C., Aratyn-Schaus, Y., and Waterman, C.M. (2010). Mechanical integration of actin and adhesion dynamics in cell migration. Annu. Rev. Cell Dev. Biol. 26, 315–333.

Gould, C.M., and Courtneidge, S. a (2014). Regulation of invadopodia by the tumor microenvironment. Cell Adh. Migr. 8, 1–10.

Haage, A., and Schneider, I.C. (2014). Cellular contractility and extracellular matrix stiffness regulate matrix metalloproteinase activity in pancreatic cancer cells. FASEB J. 28, 3589–3599.

Hayashi, T., Hirshman, M.F., Kurth, E.J., Winder, W.W., and Goodyear, L.J. (1998). Evidence for 5[prime] AMP-activated protein kinase mediation of the effect of muscle contraction on glucose transport. Diabetes 47, 1369–1373.

Vander Heiden, M.G., Cantley, L.C., and Thompson, C.B. (2009). Understanding the Warburg effect: the metabolic requirements of cell proliferation. Science 324, 1029–1033.

Hirata, H., Tatsumi, H., Lim, C.T., and Sokabe, M. (2014). Force-dependent vinculin binding to talin in live cells: a crucial step in anchoring the actin cytoskeleton to focal adhesions. Am. J. Physiol. Cell Physiol. 306, C607–20.

Van Horssen, R., Buccione, R., Willemse, M., Cingir, S., Wieringa, B., and Attanasio, F. (2013). Cancer cell metabolism regulates extracellular matrix degradation by invadopodia. Eur. J. Cell Biol. 92, 113–121.

Jameson, D.M., Thomas, V., and Zhou, D.M. (1989). Time-resolved fluorescence studies on NADH bound to mitochondrial malate dehydrogenase. Biochim. Biophys. Acta 994, 187–190.

Kanchanawong, P., Shtengel, G., Pasapera, A.M., Ramko, E.B., Davidson, M.W., Hess, H.F., and Waterman, C.M. (2010). Nanoscale architecture of integrin-based cell adhesions. Nature 468, 580–584.

Kim, D.H., and Wirtz, D. (2013). Focal adhesion size uniquely predicts cell migration. FASEB J. 27, 1351–1361.

Lakowicz, J.R., Szmacinski, H., Nowaczyk, K., and Johnson, M.L. (1992). Fluorescence lifetime imaging of free and protein-bound NADH. Proc. Natl. Acad. Sci. U. S. A. 89, 1271–1275.

Levental, K.R., Yu, H., Kass, L., Lakins, J.N., Egeblad, M., Erler, J.T., Fong, S.F.T., Csiszar, K., Giaccia, A., Weninger, W., et al. (2009). Matrix crosslinking forces tumor progression by enhancing integrin signaling. Cell 139, 891–906.

Liang, E.I., Mah, E.J., Yee, A.F., and Digman, M.A. (2017). Correlation of focal adhesion assembly and disassembly with cell migration on nanotopography. Integr. Biol. 9, 145–155.

Liberti, M. V., and Locasale, J.W. (2016). The Warburg Effect: How Does it Benefit Cancer Cells? Trends Biochem. Sci. 41, 211–218.

Lien, E.C., Lyssiotis, C.A., and Cantley, L.C. (2016). Metabolic Reprogramming by the PI3K-Akt-mTOR Pathway in Cancer. In Recent Results in Cancer Research.

Fortschritte Der Krebsforschung. Progres Dans Les Recherches Sur Le Cancer, (Springer, Cham), pp. 39–72.

Liu, J., Wang, Y., Goh, W.I., Goh, H., Baird, M. a., Ruehland, S., Teo, S., Bate, N., Critchley, D.R., Davidson, M.W., et al. (2015). Talin determines the nanoscale architecture of focal adhesions. Proc. Natl. Acad. Sci. 201512025.

Ma, N., Digman, M.A., Malacrida, L., and Gratton, E. (2016). Measurements of absolute concentrations of NADH in cells using the phasor FLIM method. Biomed. Opt. Express 7, 2441–2452.

Martin, K., Reimann, A., Fritz, R.D., Ryu, H., Jeon, N.L., and Pertz, O. (2016). Spatio-temporal co-ordination of RhoA, Rac1 and Cdc42 activation during prototypical edge protrusion and retraction dynamics. Sci. Rep. 6, 21901.

Massia, S.P., and Hubbell, J. a (1991). Human endothelial cell interactions with surface-coupled adhesion peptides on a nonadhesive glass substrate and two polymeric biomaterials. J. Biomed. Mater. Res. 25, 223–242.

Morris, B.A., Burkel, B., Ponik, S.M., Fan, J., Condeelis, J.S., Aguire-Ghiso, J.A., Castracane, J., Denu, J.M., and Keely, P.J. (2016). Collagen Matrix Density Drives the Metabolic Shift in Breast Cancer Cells. EBioMedicine 13, 146–156.

Ochsner, M., Textor, M., Vogel, V., and Smith, M.L. (2010). Dimensionality Controls Cytoskeleton Assembly and Metabolism of Fibroblast Cells in Response to Rigidity and Shape. PLoS One 5, e9445.

Palorini, R., Simonetto, T., Cirulli, C., and Chiaradonna, F. (2013). Mitochondrial complex I inhibitors and forced oxidative phosphorylation synergize in inducing cancer cell death. Int. J. Cell Biol. 2013, 243876.

Paszek, M.J., Zahir, N., Johnson, K.R., Lakins, J.N., Rozenberg, G.I., Gefen, A., Reinhart-King, C.A., Margulies, S.S., Dembo, M., Boettiger, D., et al. (2005). Tensional homeostasis and the malignant phenotype. Cancer Cell 8, 241–254.

Petit, V., and Thiery, J.P. (2000). Focal adhesions: structure and dynamics. Biol. Cell 92, 477–494.

Ponti, a, Machacek, M., Gupton, S.L., Waterman-Storer, C.M., and Danuser, G. (2004). Two distinct actin networks drive the protrusion of migrating cells. Science 305, 1782–1786.

Provenzano, P.P., Inman, D.R., Eliceiri, K.W., Knittel, J.G., Yan, L., Rueden, C.T., White, J.G., and Keely, P.J. (2008). Collagen density promotes mammary tumor initiation and progression. BMC Med. 6, 11.

Provenzano, P.P., Eliceiri, K.W., and Keely, P.J. (2009). Multiphoton microscopy and fluorescence lifetime imaging microscopy (FLIM) to monitor metastasis and the tumor microenvironment. Clin. Exp. Metastasis 26, 357–370.

Raub, C.B., Suresh, V., Krasieva, T., Lyubovitsky, J., Mih, J.D., Putnam, A.J., Tromberg, B.J., and George, S.C. (2007). Noninvasive assessment of collagen gel microstructure and mechanics using multiphoton microscopy. Biophys. J. 92, 2212–2222.

Raub, C.B., Unruh, J., Suresh, V., Krasieva, T., Lindmo, T., Gratton, E., Tromberg, B.J., and George, S.C. (2008). Image correlation spectroscopy of multiphoton images correlates with collagen mechanical properties. Biophys. J. 94, 2361–2373.

Rossow, M.J., Sasaki, J.M., Digman, M.A., and Gratton, E. (2010). Raster image correlation spectroscopy in live cells. Nat. Protoc. 5, 1761–1774.

Sameni, S., Syed, A., Marsh, J.L., and Digman, M.A. (2016). The phasor-FLIM fingerprints reveal shifts from OXPHOS to enhanced glycolysis in Huntington Disease. Sci. Rep. 6, 34755.

Schindelin, J., Arganda-Carreras, I., Frise, E., Kaynig, V., Longair, M., Pietzsch, T., Preibisch, S., Rueden, C., Saalfeld, S., Schmid, B., et al. (2012). Fiji: an open-source platform for biological-image analysis. Nat. Methods 9, 676–682.

Scott, K.E.N., Wheeler, F.B., Davis, A.L., Thomas, M.J., Ntambi, J.M., Seals, D.F., and Kridel, S.J. (2012). Metabolic regulation of invadopodia and invasion by acetyl-CoA carboxylase 1 and de novo lipogenesis. PLoS One 7, e29761.

Seewaldt, V. (2014a). ECM stiffness paves the way for tumor cells. Nat. Med. 20, 332–333.

Seewaldt, V. (2014b). ECM stiffness paves the way for tumor cells. Nat. Med. 20, 332–333.

Skala, M.C., Riching, K.M., Bird, D.K., Gendron-Fitzpatrick, A., Eickhoff, J., Eliceiri, K.W., Keely, P.J., and Ramanujam, N. (2007). In vivo multiphoton fluorescence lifetime imaging of protein-bound and free nicotinamide adenine dinucleotide in normal and precancerous epithelia. J. Biomed. Opt. 12, 24014.

Stringari, C., Edwards, R.A., Pate, K.T., Waterman, M.L., Donovan, P.J., and Gratton, E. (2012). Metabolic trajectory of cellular differentiation in small intestine by Phasor Fluorescence Lifetime Microscopy of NADH. Sci. Rep. 2.

Tilghman, R.W., Blais, E.M., Cowan, C.R., Sherman, N.E., Grigera, P.R., Jeffery, E.D., Fox, J.W., Blackman, B.R., Tschumperlin, D.J., Papin, J.A., et al. (2012). Matrix Rigidity Regulates Cancer Cell Growth by Modulating Cellular Metabolism and Protein Synthesis. PLoS One 7, e37231.

Warburg, O., Wind, F., and Negelein, E. (1927). THE METABOLISM OF TUMORS IN THE BODY. J. Gen. Physiol. 8, 519–530.

Ward, P.S., and Thompson, C.B. (2012). Metabolic reprogramming: a cancer hallmark even warburg did not anticipate. Cancer Cell 21, 297–308.

Wells, R.G. (2008). The role of matrix stiffness in regulating cell behavior. Hepatology 47, 1394–1400.

Wright, B.K., Andrews, L.M., Jones, M.R., Stringari, C., Digman, M.A., and Gratton, E. (2012). Phasor-FLIM analysis of NADH distribution and localization in the nucleus of live progenitor myoblast cells. Microsc. Res. Tech. 75, 1717–1722.

Yang, L., Hou, Y., Yuan, J., Tang, S., Zhang, H., Zhu, Q., Du, Y., Zhou, M., Wen, S., Xu, L., et al. (2015). Twist promotes reprogramming of glucose metabolism in breast cancer cells through PI3K/AKT and p53 signaling pathways. Oncotarget 6, 25755–25769.

Zhao, J., Zhang, J., Yu, M., Xie, Y., Huang, Y., Wolff, D.W., Abel, P.W., and Tu, Y. (2013). Mitochondrial dynamics regulates migration and invasion of breast cancer cells. Oncogene 32, 4814–4824.

